# The evolution of collective infectious units in viruses

**DOI:** 10.1101/524694

**Authors:** Asher Leeks, Rafael Sanjuán, Stuart A. West

## Abstract

Viruses frequently spread among cells or hosts in groups, with multiple viral genomes inside the same infectious unit. These collective infectious units can consist of multiple viral genomes inside the same virion, or multiple virions inside a larger structure such as a vesicle. Collective infectious units deliver multiple viral genomes to the same cell simultaneously, which can have important implications for viral pathogenesis, antiviral resistance, and social evolution. However, little is known about why some viruses transmit in collective infectious units, whereas others do not. We used a simple evolutionary approach to model the potential costs and benefits of transmitting in a collective infectious unit. We found that collective infectious units could be favoured if cells infected by multiple viral genomes were significantly more productive than cells infected by just one viral genome, and especially if there were also efficiency benefits to packaging multiple viral genomes inside the same infectious unit. We also found that if some viral sequences are defective, then collective infectious units could evolve to become very large, but that if these defective sequences interfered with wild-type virus replication, then collective infectious units were disfavoured.

## Introduction

Viruses disperse from host cells in many different ways. Some viruses disperse in single virions which each contain one genome. Other viruses can disperse in groups, with multiple genomes in the same virion, or multiple virions inside a larger structure. These are called collective infectious units (CIUs), and are characterised by multiple viral genomes transmitting as part of the same infective structure (Sanjuán 2017). The simplest collective infectious units consist of virions containing multiple genomes, and in viruses such as ebolavirus and paramyxoviruses, these polyploid virions can contain a variable number of genome copies (Luque et al. 2009; Rager et al. 2002). In other cases, collective infectious units can comprise larger structures containing multiple virions. These can form through free virions aggregating after dispersal, either through direct contact with one another or through collectively binding to a vector, such as a bacterial cell (Bald & Briggs 1937; Cuevas et al. 2017; Erickson et al. 2018). Alternatively, multiple virions can collectively disperse from the same host cell, for example inside extracellular vesicles formed of sections of host cell membrane, or inside protein-coated occlusion bodies (Altan-Bonnet & Chen 2015; Chen et al. 2015; Santiana et al. 2018; Slack & Arif 2007). These various kinds of collective infectious units appear to have evolved independently many times in many different viral families.

Transmitting as part of a CIU can have important consequences for viral evolution. By allowing the same host cell to be infected by multiple viral genomes simultaneously, CIUs allow for interactions between viruses even when we would otherwise expect coinfection to be rare, such as when there are strong population bottlenecks or low ratios of infectious viral particles to susceptible host cells (McCrone & Lauring 2018; Sanjuán 2018). Interactions between viral sequences can have important consequences for viral pathogenesis, diversity, and the evolution of antiviral resistance (Bordería et al. 2015; Leeks et al. 2018; Tanner et al. 2014; Vignuzzi et al. 2006; Xue et al. 2016). Furthermore, CIUs allow for repeated interactions between viral sequences, and this sets the stage for viral social adaptations. This can include cooperation, where viruses evolve adaptations that benefit other viruses, but may more commonly facilitate conflict, as in the case of defective interfering (DI) genomes, which exploit the cellular machinery of coinfecting viruses (Chao & Elena 2017; Díaz-Muñoz et al. 2017; Huang & Baltimore 1970; Sanjuán 2018; Turner & Chao 1999).

A number of hypotheses have been proposed for why viruses might transmit in CIUs. One possibility is that cells infected by multiple viral genomes might lead to more productive infections than cells infected by just one viral genome (Andreu-Moreno & Sanjuán 2018; Borges et al. 2018; Guo et al. 2017; Landsberger et al. 2018; Stiefel et al. 2012; Xue et al. 2016). In this case, CIUs might evolve if they are an effective way of delivering multiple viral genomes to the same cell (Sanjuán 2017). A second mechanism could be if CIUs allow for more efficient use of limited resources, and therefore viruses could evolve larger burst sizes by packaging multiple genomes into the same infectious unit. A third mechanism could be if viruses have a high likelihood of producing defective genomes. In that case, CIUs could be favoured to ensure that at least one functional copy of each gene is delivered to a cell, or to increase the chance that one or more complete genomes arrive in a host cell (Andino & Domingo 2015; Stiefel et al. 2012).

We model the theoretical plausibility of these three types of hypothesis: (i) if cells infected by multiple viral genomes are more productive (group infection benefits); (ii) if packaging multiple genomes into the same unit is more efficient (efficiency benefits); (iii) if there is a high likelihood that genomes are defective (insurance benefits). We ask whether each kind of hypothesis can plausibly favour the evolution of collective infectious units. For each case, we investigate what conditions are required for CIUs to be favoured as well as what sizes of CIU are favoured. Do we expect to see CIUs in all viruses, most viruses, or only under special conditions? Are some kinds of viruses more likely to evolve CIUs than others? And when CIUs do evolve, do we expect them to be small, containing just a few viral genomes, or large?

## Model

Our goal is to examine the general theoretical plausibility of potential mechanisms, rather than to capture the specific details of a single species. We have therefore purposefully left out a number of potentially important details, such as complementation between defective mutants and beneficial interactions between different variants (Andino & Domingo 2015; Leeks et al. 2018; Xue et al. 2016). We have chosen to model hypotheses which could apply to many viruses, and which could vary in predictable ways. Furthermore, we have focused on modelling the number of genomes that a generic infectious unit should contain, where an infectious unit is any structure that can deliver viral genomes to new host cells. Therefore, infectious units could reflect different biological structures, including virions, extracellular vesicles, or occlusion bodies. Our aim is to generate testable predictions across a range of different CIUs and consequently to encourage interplay between theory and data in the study of collective infectious units.

### Model Lifecycle

We imagine an acute, lytic virus spreading within a host. We assume that natural selection acts in order to maximise the rate at which it spreads. We therefore define a viral genotype’s fitness as equivalent to the expected number of future infected cells from a given infected cell. We assume that superinfection is rare enough to be ignored and that the viral progeny which leave a cell are identical to the viral genotype which initially infected the cell. Consequently, we express viral fitness, W, as:

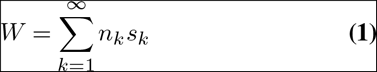

Where *k* is the number of genomes inside each infectious unit, *n*_*k*_ is the number of infectious units of size *k* produced and *s*_*k*_ is the expected number of future cellular infections each of these virions will lead to, scaled between 0 and 1. Next, we simplify our fitness equation so that we can compare the fitness of viral variants that transmit in infectious units of different sizes *k*:

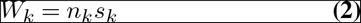

We assume that the number of infectious units that can be produced per unit time (*n*_*k*_) depends on both the number of viral genomes produced in the cell and the number of genomes that are packaged into each infectious unit. The total number of viral genomes produced by a virus may depend on the size of the infectious unit, because viruses with larger infectious units may use gene products more efficiently and so produce more genomes (see section 3.3: efficiency benefits). Consequently, we arrive at our general fitness equation:

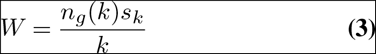

Where *n*_*g*_(*k*) is the number of genomes produced by a virus which disperses in infectious units of size *k*.

Equation 3 reveals that there is a trade-off between the number of infectious units that can be produced and the number of genomes inside each infectious unit (Fig. 1a). This tradeoff is analogous to that between the number and size of offspring (clutch size) produced by animals: with all other factors equal, the larger the clutch size, the fewer clutches can be produced (Godfray et al. 1991; Lack 1947). We will now consider three factors that could potentially favour CIUs.

**Fig. 1.**
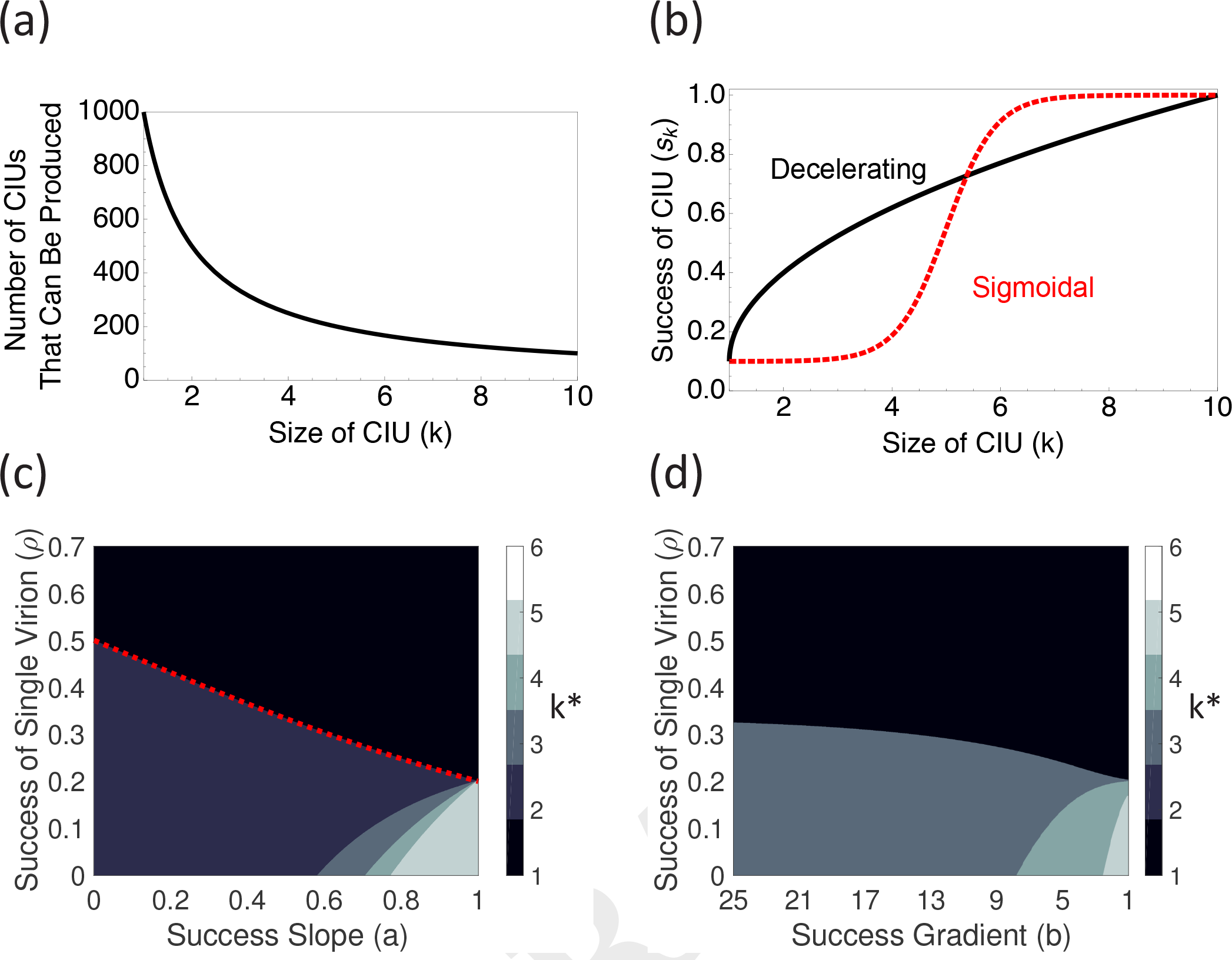
Group infection benefits and CIU evolution. (a) plots the opportunity cost of larger CIUs. All else being equal, fewer CIUs can be produced if each CIU contains more genomes. In our model, we only use integer values of *k*. (b) plots the relationship between the success of a CIU and the number of genomes it contains. (c) and (d) plot the optimal size of a CIU (*k*\*) when CIU success has a diminishing (c) or threshold (d) relationship with CIU size. The red dashed line in (c) plots the analytical condition for when CIUs evolve. When infectious unit success has diminishing returns (c), larger CIUs (*k*\* > 2) only evolve when the success slope is relatively flat (*a* is high). In contrast, when there are threshold effects (d), only larger CIUs (*k^∗^* > 2) are found, but these are found over less of the parameter space.

### Group Infection Benefits

We first consider the possibility that infections initiated with multiple viral genomes are more successful. We assume that the expected number of future infections is larger for infectious units containing more genomes, by making *s*(*k*) an increasing function of *k*. This could capture different biological mechanisms, including: larger infectious units lasting longer in the environment and so surviving longer to infect a host cell; larger numbers of initial genomes leading to a faster rate of viral production throughout the course of the cellular infection; larger infectious units having a greater likelihood of initially establishing an infection, for example through overcoming cellular immune responses, or if stochastic events early in infection can cause infections to fail (Andreu-Moreno & Sanjuán 2018; Stiefel et al. 2012). The benefit to infectious units with more genomes is analogous to when animals experience benefits through dispersing in groups, rather than alone (Davies et al. 2012; Hamilton 1971).

We assume that there is a limit to the potential benefit of multiple viral genomes infecting the same cell, and consequently that beyond a certain number of genomes, defined as *k*_*t*_, additional genomes no longer increase the productivity of an infected cell. Since we are interested in the relative fitness of different infectious unit sizes, we set the maximum potential benefit of larger CIUs, which is found at *k*_*t*_, equal to 1 and we express the success of infectious units of different sizes relative to this maximum potential benefit (y-axis of Fig. 1b). We also assume that the number of viral genomes produced per infected cell is constant, and that there is consequently a linear trade-off between the number of infectious units that can be produced and the number of genomes in each infectious unit (Fig. 1a). We consider the cases where the relationship between the number of viral genomes (*k*) and the productivity of an infected cell (*s_k_*) either shows diminishing returns, or a threshold effect (Fig. 1b; Appendix 1).

We are interested both in when CIUs evolve, and in the size of CIUs that evolve. We therefore search for the size of infectious unit (*k*) that maximises viral fitness as defined in Equation 3. We denote this value *k*\*, and it represents a candidate evolutionarily stable strategy, meaning that it could not be outcompeted by a virus employing any other strategy (Maynard Smith & Price 1973). When *k*\* > 1, CIUs are favoured over individual transmission. To find *k*\*, we evaluate our fitness equation (Equation 3) numerically at a large number of different values of *k* to determine the value which results in the highest fitness (Fig. 1 c-d). In section 2 of the Appendix, we derive an analytical condition for when collective transmission can be favoured over individual transmission when the group benefit shows diminishing returns, which we overlay in Fig. 1c.

We found that CIUs were more likely to evolve when: (i) infections initiated by a single viral genome are relatively unsuccessful (low *ρ*) (Fig. 1c-d); (ii) a small number of initial infecting genomes can reach the maximal infection efficiency (low *k*_*t*_); (iii) additional genomes have a greater influence on infection success when there are fewer genomes infecting a cell (a steeper success gradient; Fig. 1b-d); (iv) additional genomes result in a diminishing relationship with infection success (Fig. 1b-d).

We found that the conditions that favoured large CIUs were not the same as those that favoured CIUs per se. In particular, larger CIUs were favoured when: (i) infections initiated by a single viral genome are relatively unsuccessful (low) (Fig. 1c-d); (ii) a large number of initial infecting genomes are required for a successful infection (high *k*_*t*_); (iii) additional genomes have a constant influence on infection success (a shallower success gradient; Fig. 1b-d); (iv) additional genomes show a threshold effect, resulting in a sigmoidal re lationship with infection success (Fig. 1b-d).

For many factors (ii-iv above), we found that conditions that allowed CIUs to evolve more easily also favoured the evolution of smaller CIUs (Table 1). This pattern occurred because viruses are able to produce more CIUs if those CIUs are smaller (Fig. 1a). Consequently, CIUs were more likely to evolve when smaller CIUs were more successful, since in these cases viruses could achieve both the advantages of collective benefit and the advantages of transmitting large numbers of infectious units. In contrast, when the advantages of collective benefit were only possible with large numbers of genomes, viruses were able to transmit fewer of these collective units, and so CIUs were less likely to evolve.

### Efficiency Benefits

A second hypothesis for the evolution of CIUs is that they may allow a more efficient way of packaging genomes into infectious units. There are two ways that efficiency benefits could result in increased viral fitness. The first way is that there could be a limited number of structures available for collective transmission, and so packaging more genomes inside each structure could allow for more genomes to be transmitted via the collective route. This mechanism assumes either that more viral genomes are produced than can be transmitted (if CIUs are essential for transmission), or that there is an intrinsic benefit to transmitting in a CIU as opposed to transmitting as an individual virion (if CIUs are non-essential for transmission). However, there seemed no reason to assume that more viral genomes are produced than transmitted, and the second condition requires that there is already a benefit to transmitting collectively.

Therefore, we instead focus on a second hypothesis for efficiency benefits, that viruses with more efficient packaging can evolve to produce higher numbers of genomes. This hypothesis requires two assumptions. First, that larger infectious units are more efficient at packaging genomes than smaller infectious units. This occurs when a single CIU containing multiple genomes costs fewer resources than the equivalent number of infectious units containing one genome. A general way that this could occur is if the resources required to produce an infectious unit increase with surface area, and the number of genomes that it can carry depends on its volume, in which case the potential efficiency gains depend on the ratio of volume to surface area as the infectious unit increases in size. Therefore, potential efficiency gains will be greatest in infectious units which are more spherical, and which enlarge by lengthening in all dimensions simultaneously, rather than by lengthening just in one dimension.

The second requirement for this hypothesis is that the increased efficiency benefits allow for a greater number of viral genomes to be produced. One way in which this might occur is if the infectious unit is constructed from virus-derived gene products, such as structural proteins, as occurs with polyploid virions and baculovirus occlusion bodies. In this case, a more efficient use of viral proteins would result in fewer viral proteins being required to transmit the same number of viral genomes. Since viral genome copies and viral genome products are both produced by transcription of the viral genome, viruses which use these structural proteins more efficiently could evolve to produce more viral genomes (Chao & Elena 2017).

We found that the greatest efficiency benefits occurred when infectious units were spherical and when more efficient infectious units allowed more genomes to be produced. In that case, efficiency benefits scaled with the cubic root of *k* (Fig. 2a; Appendix 3). These maximum efficiency benefits therefore increased more slowly than the cost of including additional genomes (Fig. 1a), and so efficiency benefits alone were not able to favour the evolution of collective infectious units.

**Fig. 2.**
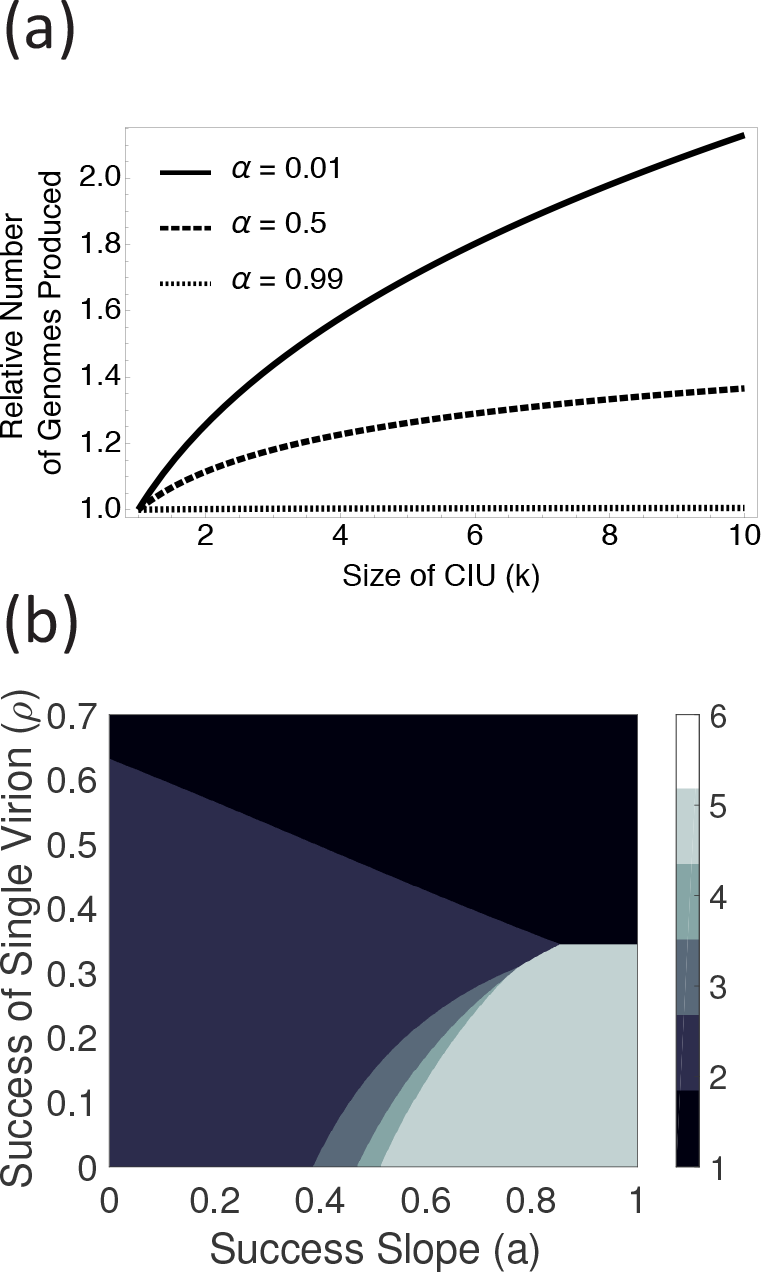
The influence of efficiency benefits on CIU evolution. (a) plots the potential increase in genome availability which comes from transmitting in CIUs of larger sizes. The increased genome availability depends on *α*, which reflects the extent to which increased efficiency of genome packaging results in more viral genome copies being produced. (b) plots the optimal size of CIU (*k*\*) which is reached for a spherical CIU with *α* = 0, reflecting the largest possible efficiency gains from larger CIUs. Compared to Fig. 1c, where there are no efficiency gains, CIUS evolve in a larger region of parameter space and are larger when they do evolve.

However, we did find that efficiency benefits were able to favour CIUs, and lead to larger CIUs, if combined with group infection benefits (Fig. 2b; section 3.2). This suggests that the requirements for CIUs to be favoured by group infection benefits may be lower when there are greater potential efficiency gains from CIUs. We found that this result critically depended on the assumption that more efficient infectious units could result in more genomes being produced (Fig. 2b).

### Defective and Defective Interfering Genomes

The third hypothesis we investigate rests on the fact that viral replication is error prone, and so some proportion of viral progeny are defective, meaning that they lack functional copies of genes required for successful infection. A high error rate could favour the evolution of CIUs, since a larger infectious unit may have a greater likelihood of containing at least one functional genome. However, in most viruses, some fraction of defective genomes are also interfering, meaning that they reduce the accumulation of the wild-type, for example if they are preferentially replicated at the expense of the wild-type genome (Huang & Baltimore 1970; Jaworski & Routh 2017; Manzoni & López 2018; Rezelj et al. 2018). Defective interfering genomes may disfavour the evolution of CIUs since larger infectious units may be more likely to contain an interfering genome. Here we incorporate both defective genomes and defective interfering genomes to see how these factors influence CIU evolution.

We investigate the possibility for defective genomes by assuming that a proportion *μ* of genomes produced are defective. For mathematical simplicity, we assume that these defective genomes are unable to be replicated in infected cells, and consequently that they don’t contribute to the success of infectious units. Therefore, this model captures the idea that collective infection could make it more likely that at least one complete genome infects a host cell (’insurance benefits’), but this model does not allow for defective genes to be trans-complemented by functional copies of the same gene in a different genome (’trans-complementation’) (Andino & Domingo 2015; Stiefel et al. 2012). By ignoring transcomplementation, our model may over-or underestimate the cost of defective genomes, since we do not allow defective genomes to contribute to group infection benefits, but we also do not allow defective genomes to build up over multiple generations. However, incorporating these additional complexities within our model would require a different model structure, since the model would need to track different classes of virus over multiple generations.

**Table 1.** Summary of theoretical predictions.

| Class of Benefit | Factor | Effect on Likelihood of CIU | Effect on Size of CIU |
| --- | --- | --- | --- |
| Group Infection Benefits | Diminishing returns (decelerating relationship) | More likely | Smaller |
|  | Threshold effect (sigmoidal relationship) | Less likely | Larger |
|  | Steeper relationship | More likely | Smaller |
|  | Low success rate of individual virion | More likely | Larger |
|  | High threshold number of genomes | Less likely | Larger |
| Efficiency Benefits | Larger CIU transmits genomes more efficiently than multiple smaller CIUs | More likely | Larger |
|  | More efficient use of CIU allows viral genome to be replicated more | More likely | Larger |
| Insurance Benefits | Lots of defective sequences | No effect | Larger |
|  | Lots of defective interfering sequences | Less likely | Smaller |

We found that defective genomes did not make CIUs more likely to evolve, but that they did influence the size of CIU that evolved (Fig. 3c; Appendix 4). When there was a very high likelihood of progeny genomes being defective (high *μ*), CIUs could be favoured to become very large, up to a second threshold, *k*_*t*_′, which is given by the value of *k* at which the likelihood of containing at least *k*_*t*_ complete genomes is approximately 1 (Fig. 3a).

**Fig. 3.**
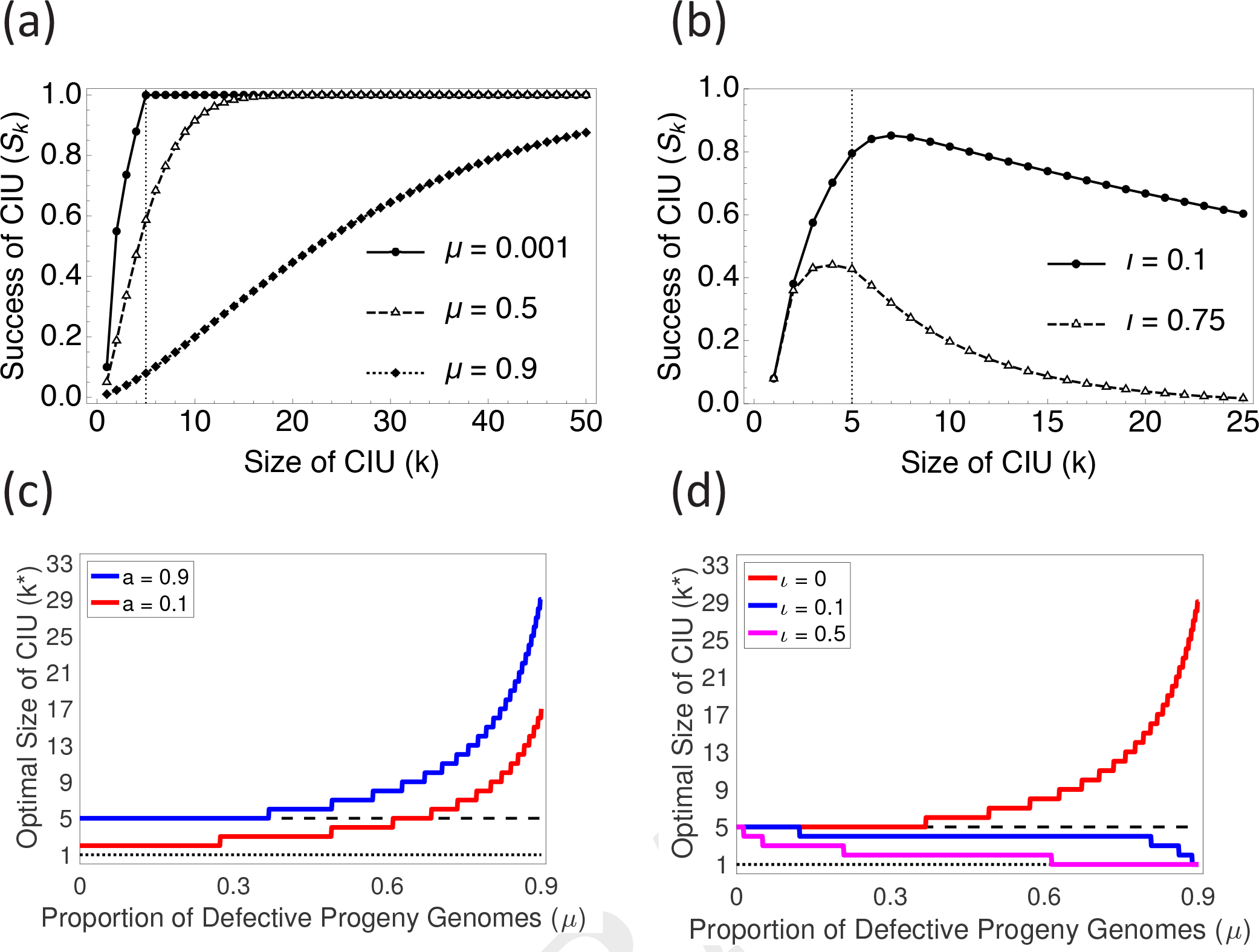
Defective and interfering genomes and CIU evolution. (a) and (b) plot the relationship between the success of an infectious unit and its size when the proportion of genomes which are defective (*μ*) (a) or which are defective and interfering (*ι*) (b) varies. In (a), as *μ* increases, virions need to be larger to achieve the same success, because there is a larger chance that the genomes inside a virion are defective. However, when some defective genomes are interfering (*ι* > 0) (b), there is a cost to larger CIUs, because larger CIUs have a greater chance of including an interfering genome. This cost reduces both the value of *k* at which success peaks and the success experienced by an infectious unit containing *k* genomes. In (b), 25% of viral progeny are defective (*μ* = 0.25). (c) and (d) plot the optimal size of infectious unit (*k*\*) as the proportion of defective genomes (*μ*) increases. The dashed line plots *k*_*t*_ (the number of complete genomes that results in maximum infectious unit success) and the dotted line plots *k*\* = 1 (when CIUs are not favoured). In (c), *a* is the shape parameter for the diminishing returns success curve, with higher values indicating a more linear curve. As defective genomes become more prevalent, the optimal size of CIU increases and can reach values which are substantially higher than *k*_*t*_. However, increases in *μ* by themselves cannot drive the evolution of CIUs from no CIUs. In (d), higher values of *ι* indicate that a higher proportion of defective genomes are interfering. As the proportion of interfering genomes (*ι*) increases, the optimal size of CIU decreases, and the likelihood that CIUs are favoured at all also decreases. Interfering genomes (*ι*) have a larger impact on CIU evolution when defective genomes are common (high *μ*).

Next, we investigated the consequences of interference by assuming that a fraction *ι* of defective genomes are also interfering. For mathematical simplicity, we assume that defective interfering genomes are completely interfering, such that a cell infected by at least one defective interfering genome produces only defective interfering genomes (Kirkwood & Bangham 1994). This scenario represents an extreme case, but it allows us to capture the qualitative influence of defective interfering genomes while keeping our model tractable (Appendix 4) (Cole & Baltimore 1973).

In contrast to our findings for defective genomes, we found that interfering genomes both: (i) made CIUs less likely to evolve, and (ii) decreased the size of the CIU which evolved when CIUs were favoured (Fig. 3d). This is because larger infectious units have a greater likelihood of incorporating a defective interfering genome, which then outcompetes the wild-type virus. In our model, the cost of defective interfering genomes depended on the product of the rate of defective mutant production (*μ*) and the chance that each defective genome is interfering (*ι*; Fig. 3c). Therefore, CIUs could only be favoured in viruses that had high rates of defective genome production if there was also a very low chance that these defective genomes were interfering (Fig. 3d).

## Discussion

We tested the theoretical plausibility of three mechanisms that could favour the evolution of group dispersal in viruses inside collective infectious units (CIUs). Our models confirmed the hypothesis that if a greater number of complete viral genomes lead to more productive infections (group infection benefits), then CIUs could be favoured (Fig. 1). However, in contrast to predictions from verbal arguments, we found that: (1) the conditions which select for CIUs tend to favour smaller CIUs rather than larger ones (Table 1); (2) in the absence of group infection benefits, neither the production of defective viruses, nor more efficient packaging of genomes, favour the evolution of CIUs (Fig. 2 & 3). Furthermore, if some fraction of progeny sequences are defective interfering genomes, then this disfavours the evolution of CIUs (Fig. 3). More generally, our results illustrate that by forcing assumptions to be made explicit, formal theoretical models can lead to different predictions than those made by simple verbal arguments.

### Predictions and Data

Our ‘group infection benefits’ model suggested that CIUs should be favoured when cells infected with multiple copies of the same viral genome lead to more productive viral infections (Fig. 1). At least two experimental studies have directly investigated group infection benefits in different viruses, in vaccinia virus (VACV) and vesicular stomatitis virus (VSV) (Andreu-Moreno & Sanjuán 2018; Stiefel et al. 2012). These studies suggest that at least two mechanisms can lead to group infection benefits: (i) if multiple genome copies are able to overwhelm cellular immunity responses; (ii) if stochastic events can prevent key viral gene products being expressed early in infection. One of these studies found a sigmoidal relationship between infectious unit size and infection success (threshold effects), and in both studies, the benefits of collective infection were large, increased relatively quickly, and saturated at relatively low numbers of genomes (*k*_*t*_ ≈ 3 genomes in Andreu-Moreno & Sanjuán; *k*_*t*_ ≈ 8 genomes in Stiefel et al.). If these studies are representative, then group infection benefits to infection could provide a relatively general explanation for the evolution of collective infectious units (Fig. 1).

Our ‘efficiency benefits’ model predicts that polyploid virions may evolve more readily in isometric viruses than in rod-shaped viruses. This is because isometric viruses, which have approximately spherical virions, may transmit multiple genomes more efficiently than rod-shaped virions. Furthermore, since polyploid virions are derived from virus-encoded capsid proteins, this efficiency benefit could feasibly allow these viruses to evolve to produce more genome copies (Fig. 2). However, it is unclear whether this prediction is borne out by data, and there are a number of caveats that could complicate this prediction, including: smaller capsids may be more stable; smaller capsids may be required for direct cell-cell transmission; capsid size may have antigenic consequences; rod-shaped capsids may enlarge to incorporate extra genetic material more easily (Flint et al. 2015; Graw & Perelson 2016; Hull 2009; Ojosnegros et al. 2011).

To what extent can the models that we considered explain the pattern of CIUs in nature? While we found that CIUs could evolve to a range of different sizes under the models that we considered, in reality most CIUs are known to be large, containing many viral genomes. For example, baculovirus occlusion bodies are known to contain dozens of individual virions, while enterovirus vesicles are large enough to potentially contain hundreds of virions (Chen et al. 2015; Slack & Arif 2007). We found that large CIUs such as these can evolve if: (i) group infection benefits increase slowly with the number of viral genomes (Fig. 1); (ii) group infection benefits require a high threshold number of genomes to accumulate (Fig. 1); (iii) viral progeny are frequently defective, but only rarely interfering (Fig. 3).

Our models have assumed that CIUs evolve due to the benefits of collective transmission. However, an alternative possibility is that collective transmission could be a by-product of selection for infectious units that are favoured for other reasons, such as increased infectivity or particle stability (Sanjuán 2017; Santiana et al. 2018). In that case, collective infection would be a consequence, but not a cause, of the evolution of CIUs, and so different kinds of explanations would be required to explain when CIUs evolve in nature.

### Further Implications

It has been suggested that collective infectious units may evolve due to the benefits of transcomplementation between defective viral genomes (Andino & Domingo 2015; Stiefel et al. 2012). While we did not model the possibility of trans-complementation, the potential for complementation to occur is greatest when defective mutation rates are high, since in that case a large fraction of the viral population could potentially benefit from transcomplementation. However, our model predicts that high rates of defective mutation may disfavour CIUs, by increasing the rate at which defective interfering (DI) genomes are produced (Fig. 3d). Consequently, our model predicts that the conditions which allow high levels of complementation to take place may also favour DIs, and so make it harder for CIUs to evolve.

Our model further suggests coevolutionary consequences between defective infectious (DI) genomes and CIUs. One possibility is that collective infectious units could favour the evolution of DIs by increasing the rate of cellular coinfection (Sanjuán 2017). This could mean that in some cases, CIUs can evolve only temporarily, since they then favour the evolution of DIs and consequently create conditions under which CIUs are no longer favoured. An alternative possibility is that viruses which transmit collectively may have evolved mechanisms of resistance which prevent their exploitation by DIs, despite the higher level of coinfection that would otherwise favour DIs. One such mechanism of resistance could be if CIUs mainly transmit sister genomes, for example due to intracellular compartmentalisation. Alternatively, if CIUs were only used episodically, for example during transmission between hosts, then DIs may be unable to accumulate, since they would be selected against during within-host transmission, when CIUs were not used.

## ACKNOWLEDGEMENTS

The authors would like to thank Ernesto Segredo-Otero, Oliver Pybus, and Katrina Lythgoe for useful comments and discussion. A.L. was supported by funding from the Clarendon Fund, St. John’s College, Oxford, and the BBSRC (grant number BB/M011224/1). R.S. was supported by ERC Consolidator Grant 724519 (ViS-a-ViS). This preprint makes use of the Henriques Lab bioRxiv template.

## Appendices

## Appendix 1

In this section we define the two ways in which the success of a CIU can depend on the number of genomes inside it. We assume that there is a collective benefit to multiple infection, such that cells infected by more viral genomes lead to more productive viral infections. Since we assume that superinfection is rare, the cellular multiplicity of infection depends only on the number of genomes that initially infect a cell, i.e. the size of the CIU (k). We assume that this collective benefit follows one of two possible relationships: diminishing returns, according to a decelerating function given by 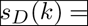 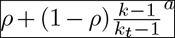; or a threshold effect, according to a sigmoidal function given by 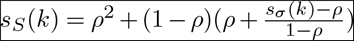 where 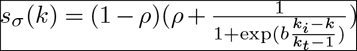. Here, *ρ* gives the relative success of a single virion, *k* is the number of genomes inside a CIU, *k*_*t*_ is the number of genomes inside a CIU at which success asymptotes, *k*_*i*_ is the value of *k* at which the inflection point of the sigmoidal curve is found, and a and b are the shape parameters for the two curves. These functions are chosen so that they always intercept the y-axis at *ρ* when *k* = 1 and at 1 when *k* = *k*_*t*_. All functions are set to have a gradient of zero when *k* > *k*_*t*_.

## Appendix 2

In this section we calculate when collective infectious units are favoured when there is collective benefit that follows a law of diminishing returns. When the success of CIUs of size *k* is given by a decelerating function, the fitness of a viral genotype producing CIUs of size *k* is given by 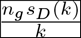 where 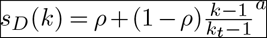 and *n*_*g*_ is a constant. We found that in this case there was always a single fitness peak with respect to size of CIU (*k*) and so CIUs evolved when the fitness of a CIU of size 2 is greater than the fitness of a CIU of size 1. We can find this by finding when *w*(2) > *w*(1), which evaluates to finding when 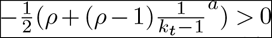. This is plotted in Fig. 2c for a given value of *k*_*t*_ and is a decreasing function of *a*, *ρ*, and *k*_*t*_, indicating that CIUs are more likely to be favoured when single virions are relatively less successful, when the benefits to additional genomes diminish rapidly, and when the benefits to collective infection can be achieved with a lower number of genomes (supplementary figure).

## Appendix 3

In this section, we derive the potential efficiency benefits that can be achieved by larger CIUs, and we calculate how these can translate into additional viral genomes. First, we assume that a CIU is spherical, since this allows for the largest possible efficiency gain. In this case, the volume of the CIU is given by 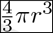 and its surface area is given by 4π*r*^2^. We assume that the number of genomes that a CIU can carry depends linearly on its volume, and that the number of constituent units that are required to build the CIU depends linearly on its surface area. We are therefore interested in the scaling relationship between the volume and surface area of the CIU; to increase the volume of the CIU by a factor of *k*, how much must its surface area increase?

We know that the volume of a CIU of size *k* is equal to *k* times the volume of a CIU of size 1. We can use this relationship to obtain an expression for the radius of a CIU of size *k*, and consequently calculate the corresponding surface area as follows. Firstly, we have that 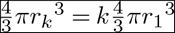 and so by rearranging, the radius of a CIU of volume *k* is 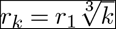 By substitution, the corresponding surface area of a CIU of volume *k* is 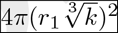. We are interested in the number of genomes which can be transmitted per unit required for constructing the CIU (‘structural units’, such as capsid proteins if the CIU is a virion). This is given by the number of genomes transmitted per CIU multiplied by the number of CIUs which can be produced from a given amount of structural unit, which we can write as 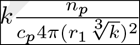 where *n*_*p*_ is a constant giving the number of structural units available, and *c*_*p*_ is a constant giving the amount of surface area of CIU that can be produced by one structural unit. We now obtain the relative efficiency advantage by dividing the number of genomes transmitted with a constant amount of structural unit available for a genotype producing CIUs of size *k* by the same expression when *k* = 1, which gives us our scaling relationship 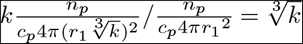.

To translate the efficiency benefit of a larger CIU into viral genome availability, we are interested in the relative number of genomes that a virus with CIUs of size *k* can produce. We first assume that the structural units required to build a CIU are virus-derived and depend on the viral genome being transcribed. We further assume that there is a linear trade-off between transcriptional events that produce viral gene products, resulting in structural units, and transcriptional events that replicate the viral genome, resulting in viral genome copies. Therefore, we assume that a reduced requirement for viral-derived structural units can result in an increased number of viral genome copies (Chao & Elena 2017). Since we are interested in the relative viral genome production, we first assume that viruses are adapted to be efficient such that when they have CIUs of size 1 (*k* = 1), genomes and CIU structural units are produced in the ratio *α* : 1 − *α*, and that at this ratio, the right number of genomes are produced to fill every CIU constructed. We now assume that when *k* > 1, a larger number of genomes can be packaged per structural unit, according to the ratio 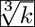 derived above. With this increased efficiency of using structural proteins, the optimal ratio of genome:structural protein production now deviates from 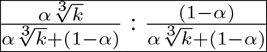 where we divide both sides of the ratio by the total amount of structural protein production and genome production, since we assume that this remains constant. Assuming that viruses with CIUs of size *k* quickly evolve to an optimal ratio of genome:structural protein production, we can express the relative number of genomes available to a virus with CIUs of size *k* by dividing the left hand side of this ratio for a general value of *k* by *α*, which yields 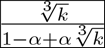 This relationship varies between 0 when *α* = 1 and 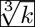 *k* when *α* = 0. This is because when *α* is low, relatively fewer genome copies are produced relative to viral gene products when CIUs are small, and so there is a larger potential gain in the number of genomes produced by viruses with larger CIUs. The relationship 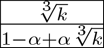 is plotted in Fig. 2a.

## Appendix 4

In this section, we calculate the impact of defective and defective interfering viral genomes on the optimal size of CIU that evolves. First, we calculate the impact of defective genomes. We assume that a fraction *μ* of progeny viral genomes are defective, and are not replicated inside an infected host cell and also do not contribute to the success of larger virions. We also assume that these defective genomes are just as likely as complete genomes to be incorporated into a CIU, and so the distribution of genomes inside CIUs is well described by a Binomial distribution, where the number of trials is the size of CIU and ‘successes’ represent incorporation of a complete genome, which happens with likelihood 1 − *μ*. The success of a CIU is now given by the sum of the product of the likelihood of each possible combinations of complete and defective genomes (each state *c* where *c* ∈ {1…*k*}), and the success of each state 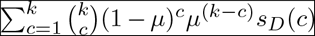 where *k* is the total number of genomes (both complete and defective) inside a CIU, *c* is the number of complete genomes inside a CIU, *μ* is the likelihood that a progeny genome is defective, and *s*_**D**_ (*c*) describes the success of a virion with *c* complete genomes. This success function is plotted in Fig. 3a and leads to a new value of *k* at which success asymptotes, *k*_*t*_′, which we can find by finding a value of *k* for which 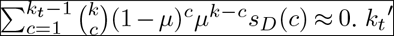 can reach very large values when *μ* is high, indicating that CIUs may evolve to become very large if defective mutations are very common.

We next incorporate the possibility that a fraction *ι* of the defective genomes are also interfering. We assume that interference is total such that any infectious unit that contains at least one interfering genome produces no wild-type genomes upon infection of a new cell. We therefore weight the likelihood of each infection state *c* by the likelihood that none of the defective genomes are interfering, and so our new expression for the success of a CIU of size *k* becomes 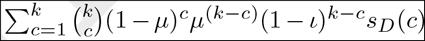, which we plot in Fig. 3b.

## Supplementary Data

MATLAB scripts for the numerical model and for reproducing figures are available at https://osf.io/v3ru8/.

